# A structural dynamic explanation for observed escape of SARS-CoV-2 BA.2 variant mutation S371L/F

**DOI:** 10.1101/2022.02.25.481957

**Authors:** Nathaniel L. Miller, Thomas Clark, Rahul Raman, Ram Sasisekharan

## Abstract

The SARS-CoV-2 Omicron sub-variants BA.1 and BA.2 have become the dominant variants worldwide due to enhanced transmissibility and immune evasion. In response to the rise of BA.1 and BA.2, two recent studies by Liu et al. and Iketani et al. provide a detailed analysis of loss of therapeutic antibody potency through evaluation of escape by pseudotyped viruses harboring BA.1 and BA.2 receptor binding domain (RBD) point mutations. Surprisingly, Liu et al. and Iketani et al. observed a profoundly broad escape effect for the individual mutations S371L and S371F. This result cannot be explained by known escape mechanisms of the SARS-CoV-2 RBD, and conflicts with existing computational and experimental escape measurements for S371 mutations performed on monomeric RBD. Through an examination of these conflicting datasets and a structural analysis of the antibodies assayed by Liu et al. and Iketani et al., we propose a mechanism to explain S371L/F escape according to a perturbation of spike trimer conformational dynamics that has not yet been described for any SARS-CoV-2 escape mutation. The proposed mechanism is relevant to Omicron and future variant surveillance as well as therapeutic antibody design.

---

Upon emergence, the SARS-CoV-2 Omicron sub-variant BA.1 was identified to have increased transmissibility^1^ and immune evasion^2^ and has since become the dominant variant worldwide. Subsequently, the Omicron sub-variant BA.2 was observed to have a growth advantage as compared to BA.1^3^. In response to the rise of BA.1 and BA.2, scientists worldwide have raced to computationally^4,5^ and experimentally^6–10^ characterize the decreased efficacy of current vaccines and therapeutic antibodies that were designed to target the wild-type Wuhan SARS-CoV-2 strain. Specifically, two recent studies by Liu et al.^9^ and Iketani et al.^10^ provide a detailed analysis of loss of potency by evaluating vaccine/convalescent sera and therapeutic antibodies against pseudotyped viruses with D614G spike proteins harboring single point mutations from the variants of concern (VOCs). This characterization of individual variant mutations improves our mechanistic understanding of receptor binding domain (RBD) antigenic space, facilitating next-generation antibody and vaccine design and interpretation of future variant phenotypes.

Surprisingly, when assaying individual BA.1 and BA.2 mutations Liu et al. and Iketani et al. observed a profoundly broad escape effect for S371L and S371F. While S371L/F mutations occur in class 4 antibody epitopes, Liu et al. and Iketani et al. observed escape from the majority of antibodies targeting all four RBD epitope classes including those on distant RBD surfaces. At the level of epitope surface spanning an individual RBD, the broad escape mediated by a mutation at single site S371 is unanticipated because this site is unlikely to have a detrimental effect across all known epitope-antibody interfaces in SARS-COV2 as defined by us^4,5^ and others^11,12^ previously. Further, S371 mutations have not been previously observed on variants of interest through the SARS-CoV-2 pandemic as would be expected if mutations at this site could produce such broad antibody escape without an associated fitness cost. We therefore sought to identify a mechanism through which isolated S371 mutations could confer broad antibody escape across all four classes of anti-RBD antibodies while also explaining the anticipated fitness tradeoff that has constrained evolution at this site prior to the emergence of Omicron.

S371L/F mediated escape is unlikely to be explained by long-range conformational changes within the protein structure of monomeric RBD. Unlike the allosteric disruption of RBD structure observed in the case of the E406W mutation^13^ that also lead to broad antibody escape, the S371 mutations did not impact ACE-2 binding and only slightly reduced monomeric RBD expression when assayed via yeast display^14^. The escape of S371L/F mutation across antibodies targeting all four epitope classes was observed only when assayed in the context of the spike trimer by Liu et al. and Iketani et al. Therefore, we structurally examined the individual antibody-antigen interactions assayed by Liu et al. and Iketani et al. in the context of the spike trimer to toward elucidating additional mechanistic details of S371L/F escape.

Investigation of class 1 to 4 antibody epitopes in the context of the open and closed states of trimeric spike suggests the S371L/F escape mechanism involves altered RBD conformational dynamics. Specifically, we mapped the epitopes of the antibodies assayed by Liu et al. and Iketani et al. to open versus closed spike protein structures and found that S371L/F-mediated escape was strongly associated with epitope accessibility in the spike closed (3 RBD_down_) vs spike open (1-3 RBD_up_) conformational states (**Figure 1**). That is, in the closed spike state class 1 and class 4 antibody epitopes are not accessible while class 2 and 3 antibody epitopes are largely accessible (Figure 1A), and the relative closed-state accessibility for each antibody predicted whether a given antibody was weakly/moderately (class 2 and 3) or strongly (class 1 and 4) escaped by S371L/F (Figure 1B).

**Figure 1:**
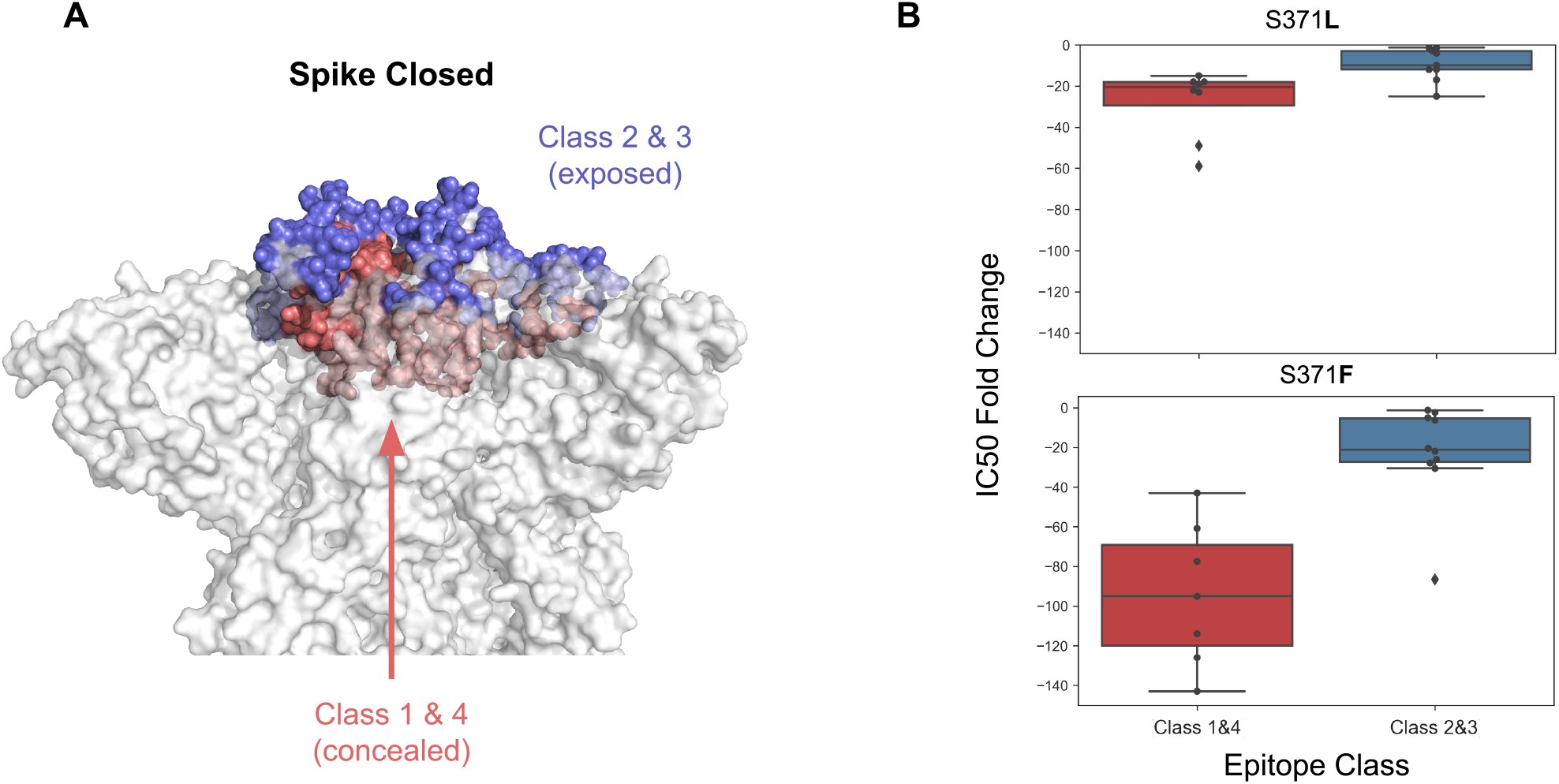
Epitope accessibility in the spike closed versus open state is associated with magnitude of S371L/F-mediated antibody escape. (A) Surface representation of spike trimer in the closed (3 RBD_down_) state (PDB: 6ZGI). Class 2 and 3 antibody epitopes are largely accessible in the spike closed state, while class 1 and 4 antibody epitopes are concealed. (B) Antibodies whose epitopes are concealed in spike closed (class 1 and 4) suffer from greater S371L/F-mediated escape than antibodies whose epitopes are largely exposed in spike-closed.

Importantly, class 2 and 3 antibody epitopes have variable antibody accessibility in the RBD-down state. To further characterize the epitope accessibility-escape trend for S371L/F and to additionally benchmark the observed trend against other BA.1 and BA.2 escape mutations, we computed the ratio of antibody epitope accessibility in the spike-closed conformation as compared to the spike-open conformation and plotted this metric against the antibody escape across BA.1 and BA.2 mutations (**Figure 2**). We found that for S371L and S371F—but not for other BA.1 or BA.1/2 escape mutations such as K417N or E484A nor for the full suite of BA.1/2 mutations—epitope *inaccessibility* in the RBD_down_ conformation was associated with greater antibody escape. Specifically, all class 1 and class 4 antibodies examined whose epitopes are not accessible in RBD_down_ were strongly escaped by S371 mutations. In contrast, class 2 and 3 antibodies whose epitopes are partially or fully accessible in RBD_down_ were escaped to a lesser extent. While the degree of class 2/3 accessibility in the closed state was correlated with escape for S371L, the association was weaker for S371F suggesting that additional features contributing to escape are at play, particularly across class 3 antibodies. The variance for class 3 antibody escape may be explained by the hypothesis offered by Liu et al. regarding modulation of the N343 glycan, as class 3 antibody epitopes are proximal to this N-glycan (5.4 Å between N-glycan and nearest REGN10987 light chain heavy atom) and in the case of S309 even involve direct contacts with this N-glycan. Additionally, REGN10987 (the largest outlier from the trend) features steric constraints on binding in the closed state resulting from clashes between the light chain and the adjacent RBD despite an epitope that is highly accessible.

**Figure 2:**
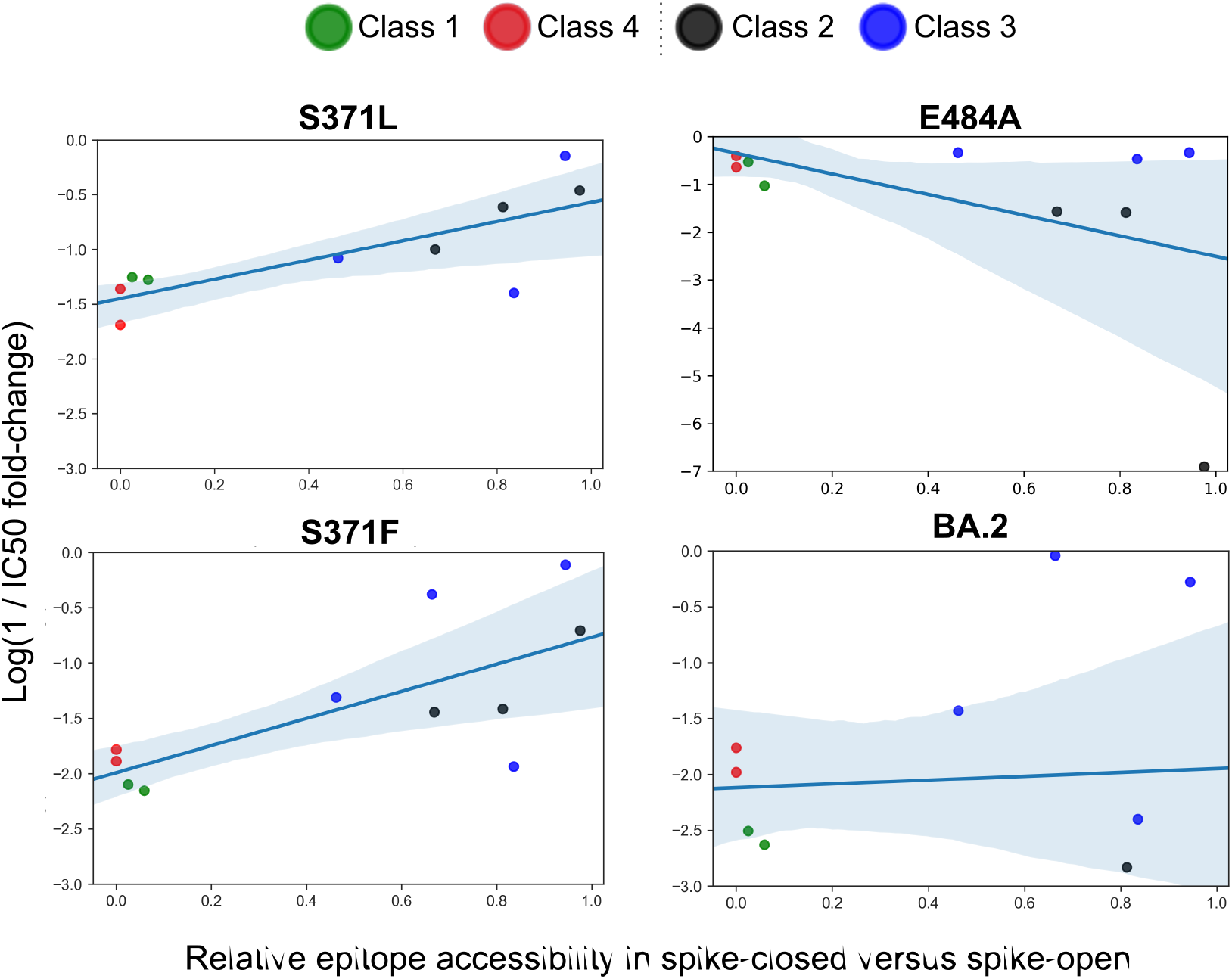
Antibody epitope accessibility in RBD_up_ versus RBD_down_ correlates with observed S371L/F escape. Escape (*y-axis*) measured by Liu et al.^9^ and Iketani et al.^10^ against S371L, S371F, K417N, or E484A for 11 antibodies with solved crystal structures is plotted versus RBD_down_ epitope accessibility (*x-axis*). Class 1-4 antibodies are annotated by color as shown in the legend at right. While epitope accessibility in the RBD_down_ conformation does not correlate with antibody escape from K417N or E484A escape, a correlation is observed for S371L and S371F mutations. Reduced RBD_down_ epitope accessibility results in the strongest escape for class 1 (green) and class 4 antibodies (red). Class 3 antibodies (blue) exhibit the greatest variance from this trend, and antibody REGN10987 is the primary outlier, as it suffers from large S371L/F escape despite high RBD_down_ epitope accessibility.

Therefore, the analyses of diverse data including RBD expression, ACE-2 binding, antibody binding in the monomeric RBD context, pseudovirus neutralization representing the trimeric context, and RBD_down_ versus RBD_up_ epitope accessibility, provides a plausible explanation for the observed escape effects of isolated S371 mutations based on RBD_up_ versus RBD_down_ conformational dynamics. The mechanism through which S371 mutations mediate this perturbation of RBD dynamics might include modulation of 1) RBD_down_-RBD_down_ interface interactions^4^, 2) the N343 glycan^9^, or 3) folding/stability^14^ of the RBD which only produces an escape effect in the trimeric context. In all three cases, the effect may bear a significant fitness cost such that isolated S371 mutations are not viable in the existing fitness landscape, explaining the lack of variants bearing S371 mutations in isolation. Significantly, both the S371L/F escape effect and fitness cost appear to be reduced in combination with other BA.1/BA.2 mutations. That is, the S371L/F escape effect exceeds the BA.1/BA.2 escape effect for a number of antibodies, and it is apparent from Omicron’s global dominance that S371L/F in combination with other mutations such as G339D, S37P, S375F, and T376A results in a fit lineage.

Of these three mechanisms, we find the most support for mechanism #2 involving the N343 glycan. Liu et al. had previously suggested that S371 mutations interact with the N343 glycan to confer escape from class 3 antibodies. Here, we hypothesize that interactions between L/F371 and the N343 glycan further result in altered spike-closed versus spike-open conformational dynamics to confer broad escape from class 1 and 4 antibodies whose epitopes are inaccessible in the spike-closed conformation. S371L/F may confer a predominantly “locked” spike closed state and reduced occupancy of the spike open state. Further supporting this hypothesis and providing additional mechanistic details, MD simulations previously identified the N343 glycan as a “glycan gate” control element that drives the spike opening process^15^. Altered closed versus open state occupancy also explains the expected S371L/F fitness cost, as a tradeoff between closed-state occupancy and infectivity was recently established via smFRET characterization of the altered conformational dynamics due to the D614G and Alpha variant mutations by Yang et al^16^.

In summary, we offer a plausible biological explanation for the profound escape conferred by S371L/F as measured by Liu et al and Iketani et al., and believe that further experimental and structural investigation is warranted. Future work to characterize the BA.1/BA.2 mutations that compensate for the S371L/F fitness cost would be particularly valuable toward understanding the Omicron evolutionary landscape, surveilling these compensatory sites for future mutation on Omicron or novel variants, and designing therapeutic antibodies targeting the compensatory sites which may be functionally-constrained to mutate. Two BA.2 mutations of particular interest as potential compensatory sites are D405N and R408S, as Sztain et al. previously identified these sites as participating in the “glycan gate”-mediated RBD opening process. Further, BA.1/BA.2 mutations S37P, S375F, T376A, and Y505H in the RBD_down_-RBD_down_ interface may modulate the Omicron spike-closed RBD packing^17^ and opening dynamics^4^ to compensate for S371L/F.

## Acknowledgements

We thank Sho Iketani, Lihong Liu, and David Ho for their correspondence and discussion. NLM was supported in part by T32 ES007020/ES/NIEHS NIH and by SMART, Singapore.

## Methods

### Antibody Epitope Accessibility Analysis

Antibodies assayed for pseudoviral neutralization in Liu et al.^9^ or Iketani et al.^10^ with publicly available structures were selected for analysis (PDBs: 7C01, 7CDI, 6XDG, 7L7E, 7KMG, 6XDG, 7L7E, 7JX3, 7MMO, 7LD1, 7RAL). For each antibody, RBD epitope residues were identified using the PyRosetta NeighborhoodResidueSelector() with a cutoff of 7 Å. Subsequently, the RBD_down_ to RBD_up_ accessibility ratio was computed as the ratio between the total solvent accessible surface area (SASA) for each antibody’s epitope residues on an RBD_down_ in the 3 RBD-closed conformation (PDB: 6ZGI) and an RBD_up_ (PDB: 6M0J). SASA was computed in PyRosetta using SasaCalc() and a 10 Å probe. The epitope accessibility score was then plotted versus the escape values measured in Liu et al.^9^ or Iketani et al.^10^ for each mutation using Seaborn regplot(). Note that the raw reported fold-change in IC_50_ was first converted to fold-change relative to wild-type (IC_50,mutant_ /IC_50,WT_) and then the natural logarithm was taken prior to plotting.

